# Reprogramming protein abundance fluctuations in single cells by degradation

**DOI:** 10.1101/260695

**Authors:** O Matalon, A Steinberg, E Sass, J Hausser, ED Levy

## Abstract

Isogenic cells living in the same environment show a natural heterogeneity associated with fluctuations in gene expression. When these fluctuations propagate through cellular regulatory networks, they can give rise to noise regulons, whereby multiple genes fluctuate in a coordinated fashion in single cells. The propagation of these fluctuations has been extensively characterized at the transcriptional level. For example, variations in transcription factor concentration induce correlated fluctuations in the abundance of target gene products. Here, we find that such noise regulons can also stem from protein degradation. We expressed pairs of yellow and red fluorescent proteins, subjected them to differential translation or degradation, and analyzed their fluctuations in single cells. While differential translation had little impact on fluctuations, protein degradation was found to be a dominant contributor. A mathematical model to decompose fluctuations arising from multiple sources of regulation revealed that cells with higher protein production capacity also exhibited higher protein degradation capacity. This association uncouples fluctuations in protein abundance from fluctuations in production rate, and can generate orthogonal noise regulons even for proteins relying on the same transcriptional program.

## Introduction

Molecular noise is ubiquitous in biological systems (1–7) and originates from two distinct sources (8–11). A first, intrinsic source stems from the stochastic nature of chemical reactions within cells. Considering proteins, intrinsic noise measures the variation of a protein’s concentration, when all cellular parameters are kept constant. The second source of noise is the extrinsic component, which corresponds to the variability of a protein’s concentration across different cell states. Isogenic cells living in the same environment indeed naturally explore a multitude of states reflected in differences in size, shape, cell cycle phase, concentrations of polymerases, ribosomes, regulation factors, *etc.* (12, 13). Such extrinsic noise is sometimes referred to as pathway noise (14, 15). Recent advances in single-cell RNA- seq have also contributed to unveiling the natural heterogeneity of cell populations and cell states, even allowing the identification of novel cell types (16–19).

Understanding the molecular bases driving cellular heterogeneity can yield fundamental insights into mechanisms of cell function and regulation (20–33). This idea was explored by Perdaza *et al.* who expressed two fluorescent proteins in a cascade and observed that fluctuations of the regulator propagated to the regulatee (20). More generally, correlated fluctuations of protein abundance have been widely observed in *S. cerevisiae* (21). In that work, the authors observed that proteins undergoing correlated fluctuations with a stress response factor were involved in stress response themselves. These correlations proved predictive of regulatory mechanisms although they were measured in unstressed cells (21). Identification of sources of extrinsic noise affecting a specific protein can thus reveal how proteins are regulated (22–30). Based on this idea, Farkash-Amar *et al*. used correlation in protein abundance and localization to identify 74 genes related to human cell motility (29). In another example, it was observed in yeast that cell-specific growth rate and stress resistance were anti-correlated, and the cellular abundance of Tsl1, a trehalose synthase component, correlated with slow-growing, stress- resistant cell states (31).

At the root of heterogeneity lies the question, how can extrinsic noise be produced? Sources of extrinsic noise have been extensively characterized at the transcriptional level (5, 15, 20, 23, 34–41), and recent advances of single cell transcriptomics by RNA-seq have contributed to consolidating that view (42, 43). Theoretically, however, any regulatory mechanism that can affect a protein’s level could also influence its noise (34), *e.g.*, by changing translation, mRNA, or protein degradation rates across cells. Importantly, recent advances in transcriptomics and proteomics methods have shown that post-transcriptional regulation greatly contributes to homeostasis of protein abundance (44, 45), a view also supported by single-cell measurements of mRNA and protein levels (44, 46). The fundamental role of post- transcriptional regulation in regulating protein levels is particularly well illustrated in a recent work, where yeast proteins were all expressed from the same constitutive promoter, but showed highly variable abundances, spanning over two orders of magnitude (47). At the functional level, post- transcriptional regulation is indeed crucial for many key cellular processes such as the cell cycle (48). More generally, entire classes of proteins can be subject to strict post-transcriptional regulation (49–51). For example, Gsponer *et al.* observed across several species that proteins rich in disordered regions are tightly regulated throughout their lifetime, from transcript synthesis to protein clearance (50).

The fact that post-transcriptional regulation mechanisms play a major role in cellular circuits prompts us to ask whether they represent a source of extrinsic noise on top of transcription. For example, if a protein requires a specific factor to be degraded, the fluctuations in the abundance of the protein will be coupled the fluctuations of the degradation factor.

To evaluate whether post-transcriptional processes can impact fluctuations of protein abundance in single cells, we compared fluctuation patterns of fluorescent proteins in presence or absence of sequence tags inducing either decreased translation rate or increased degradation. We used a two-color reporter strategy (8, 9) to quantify the extent of change in extrinsic noise caused by the sequence tags (Fig 1). We observed that decreased translation rate did not significantly impact extrinsic noise despite inducing a 3-fold reduction in protein abundance. Increased degradation, however, which was triggered by a misfolded polypeptide tag, caused a dramatic change in the pattern of cell-to-cell fluctuations.

**Fig 1.**
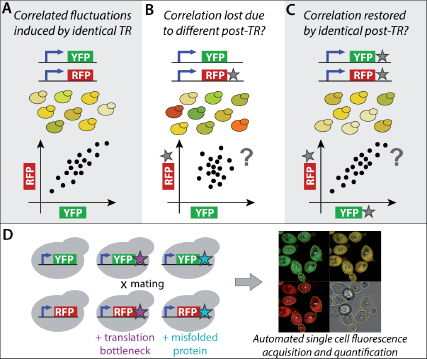
Assessing the impact of post-transcriptional regulation (post-TR) on protein abundance noise. **(A)** Two proteins, YFP and RFP are expressed under the same promoter and at the same genomic locus in diploid yeast cells. Identical promoters subject the two proteins to identical transcriptional regulation across cells, e.g., if a particular cell has a lower concentration of transcription factor, it will produce less of both YFP and RFP. Thus, correlated fluctuations of YFP and RFP across single cells are expected. **(B**) RFP* now differs from RFP and YFP by a specific sequence tag (star). If a regulator modulates RFP* abundance through that tag, *e.g.*, by degradation, then cell states with higher concentrations of the regulator will result in lower concentrations of RFP*. This new layer of regulation may confound the transcriptional layer, resulting in decreased correlation between RFP* and YFP. **(C)** Adding the same tag to YFP would restore the correlation because both proteins would once again be subjected to identical regulation. **(D)** We constructed yeast strains to assess whether post-TR can impact fluctuations of protein abundance across single cells. We used two sequence tags fused to the fluorescent proteins. The first is a sequence with low translation efficiency and the second is a truncated, misfolded protein. Combinations of these variants were expressed in yeast cells and their abundance was measured.

## Results

### Measuring protein abundance noise using a two-color reporter strategy

The noise of protein abundance in single cells can be decomposed into intrinsic and extrinsic components using two fluorescent reporter proteins of different colors, as originally proposed (8, 9). In this strategy, cell-to-cell differences impact the expression of the two reporters in the same way, such that correlation in their abundance across cells measures extrinsic noise, whereas differences in reporter abundance within cells reflect intrinsic noise.

We adopted this strategy and expressed a yellow and a red fluorescent protein in *S. cerevisiae*. The genes coding for the two reporters were integrated at homologous loci and used identical promoters (Details of constructs, strains, and sequences are given in Fig S1, Text S1 and Tables S1-S2). We measured the fluorescence of diploid yeast cells expressing the reporters using an automated confocal spinning disk microscope (Fig 1D). As expected, YFP and RFP abundance were highly correlated across cells (R=0.83, Fig 2), reflecting that both reporters were indeed influenced by identical sources of extrinsic noise.

**Figure 2.**
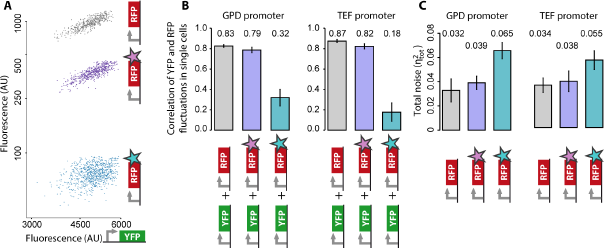
Post-transcriptional regulation can either maintain or abolish protein abundance coupling in single- cells. **(A)** The abundance of YFP (x-axis) and RFP variants (y-axis) are measured in single cells (points). We use three RFP variants: untagged RFP (grey), RFP fused to a sequence acting as a translation bottleneck (purple), and RFP fused to a misfolded protein (cyan). The YFP and RFP variants are under GPD promoter expression. **(B)** Pearson correlation between YFP and each RFP variant. Independent experiments are shown using either GPD or TEF promoters. **(C)** The total squared noise 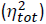 was calculated by the variance of the log-transformed fluorescence intensities (Equation 9, Methods). Error-bars in panels B and C correspond to two standard deviations calculated based on six biological replicates.

### Decreasing translation rate minimally impacts protein abundance fluctuations

We then altered the post-transcriptional regulation of only one of the reporters and analyzed the resulting impact on extrinsic noise (Fig 1B). In a first experiment, we fused an amino acid sequence at the C-terminus of RFP, which contained seven repeats of “CTT,” a leucine codon with low tRNA adaptation index in *S. cerevisiae* (52) (Methods, Text S2). We call this sequence a “translation bottleneck” (tb) and use RFP-tb to refer to this variant of RFP. As expected by design, the average cellular abundance of RFP-tb was lower than that of untagged RFP, by ~3-fold (Fig 2). Interestingly, the correlation remained close to the original value (R=0.79), indicating that fluctuations were not affected by the translation bottleneck. The total noise for RFP-tb was comparable to RFP (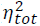 =0.039 and 0.032 respectively, Fig 2C). Overall, the similarity in correlation in presence and absence of the translation bottleneck indicates that all cells, irrespective of their state (here represented by YFP abundance), deal with the bottleneck sequence with comparable efficiency. Lastly, we repeated these experiments using a different promoter and our observations remained highly similar (Fig 2).

### A misfolded protein tag decouples protein abundance fluctuations

In a second experiment, we fused a misfolded protein to RFP (53) and refer to this variant as RFP-misP. Protein misfolding is a pervasive process that can be triggered by a stress such as heat shock (54), but can also occur during the normal life cycle of proteins due to translational errors, for example (55). As a result, cells have evolved elaborate quality-control machineries (56). By expressing RFP-misP with YFP, we tested whether the cell machinery dealing with RFP-misP would subject it to a new source of extrinsic noise, and whether that new source could decouple the fluctuations between RFP-misP and untagged YFP. We observed a 20- fold decrease in RFP-misP expression. This decrease likely originates from a change of protein stability rather than a change of mRNA stability or translation efficiency, because fusion of RFP-misP to an additional solubilizing tag rescues protein abundance (Fig S2). Strikingly, the correlation between RFP- misP and YFP underwent a large decrease, down to R=0.32. We also observed a two-fold increase in total noise (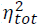 = 0.065) relative to RFP alone (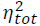 = 0.032, Fig 2C and Methods).

We performed the same measurements using a different promoter and observed similar results: the correlation between YFP and RFP-misP decreased (R=0.18), and the total noise increased due to the misP tag (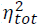 = 0.055) compared to RFP alone (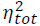 = 0.034). In another control, we swapped fluorescent reporters, using YFP-misP together with RFP, and we observed similar results. The correlation vanished (R < 0.1), and the total noise increased due to the misP tag (Fig. S3).

### Post-transcriptional co-regulation re-couples fluctuations of protein abundance in single cells

Two hypotheses could explain the decoupling of YFP and RFP-misP expression. One possibility is that tagging RFP with misP increases its intrinsic noise. In that case, tagging RFP and YFP simultaneously with misP should not re-couple fluctuations. Alternatively, RFP-misP may be subject to a new source of extrinsic noise. In this case, tagging both RFP and YFP with misP should restore the coupling in protein expression. We observed the latter scenario, where tagging both proteins with misP restored their correlation (R=0.76, Fig 3). The same experiments based on a different promoter gave similar observations (R=0.71, Fig 3C).

**Fig 3.**
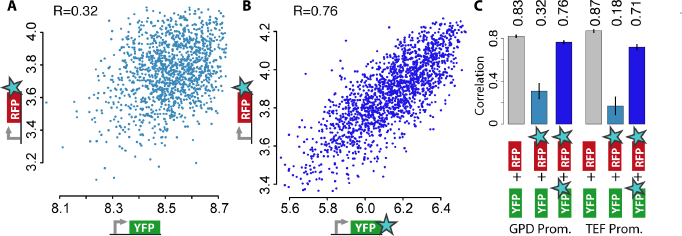
Post-transcriptional co-regulation re-couples fluctuations of protein abundance in single cells. **(A)** Fluorescence intensities measured in single cells for YFP (x-axis) and RFP-misP (y-axis), showing a weak correlation of their abundance across cells (fluorescence arbitrary units). **(B)** Expression of YFP-misP with RFP-misP restores the correlation of fluctuations across single cells. **(C)** Correlation for different pairs of proteins subjected to different post-transcriptional regulations: untagged YFP + untagged RFP (grey), untagged YFP + RFP-misP (cyan), or YFP-misP + RFP-misP (blue). Average of six biological replicates, error-bars show two standard deviations.

Thus, subjecting proteins to a new layer of post-transcriptional regulation can change their pattern of fluctuation at the single cell level. Moreover, the restoration of the correlation that we observed indicates that this change is caused by a source of extrinsic noise, supposedly reflecting the “degradation capacity” of misfolded proteins in each cell. We assessed whether this cell-specific degradation capacity could be linked to the cell cycle, and measured the correlation between the abundance of YFP-misP and RFP-misP in sub-populations of cells grouped by size or cell cycle stage. The correlation depended on neither property (Fig S4, S5), indicating that cell size and cell-cycle stage do not influence the extrinsic factor represented by the “degradation capacity.”

### Post-transcriptional regulation creates anti-fluctuations that partially cancel the transcriptional fluctuations in our system

To explain the decoupling and recoupling of protein fluctuations by post- transcriptional regulation, we introduce a mathematical model of the fluctuations of reporters subjected to multiple noise sources. Two reporters G and R sharing the same promoter and lacking the misP tag share the same source of extrinsic noise Z. Following a framework introduced by Elowitz et al. (8), we model the fluorescence of the reporters as

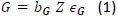

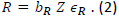

where G and R represent the single cell fluorescence of the green (GFP or YFP) and red (RFP) reporters. The *b* constants account for differences in average protein abundances and differences in abundance-to- fluorescence scaling. 𝜖 is the intrinsic noise due to stochasticity inherent to gene expression. We use a multiplicative model because, with mass action kinetics, fluctuations in protein production and degradation have multiplicative effects on protein abundance (Methods).

Adding a misfolded tag to one reporter subjects it to a new source of extrinsic noise 𝑊. In the case of RFP-misP for example, 𝑊 accounts for the effect of the misP tag on fluorescence in single cells now denoted 𝑅^∗^, with

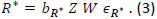

By log-transforming fluorescence (log 𝐺 → 𝐺, log 𝑅^∗^ → 𝑅^∗^) to linearize these equations, we can compute how the noise sources *Z* and *W* impact the correlation between log protein abundance of the two reporters 𝐺 and 𝑅^∗^ across a cell population by

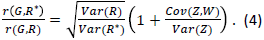

This equation (see Methods for details of the derivation) formalizes the intuition of Fig 1: subjecting *R\** to a new source of noise *W* can decrease the correlation between the reporters in two distinct mechanisms. First, *W* can inject extra noise into 𝑅^∗^ to increase *Var(R*)*. As the first term of Equation 4 shows, increasing *Var(R*)* decreases the correlation between *G* and *R\**. Second, if the coupling between *Z* and *W* is negative, fluctuations in *Z* are (partially) canceled by anti-fluctuations in *W* which decouple *R\** from *G*. This is reflected in the second term of Equation 4, where *Cov(Z,W)* << 0 decreases the correlation between *G* and *R\**.

In our experiments, 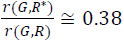 and 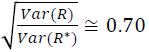, which implies *Cov(Z,W)* < 0. The data thus suggest that the loss of correlation between *G* and *R\** is due both to increased noise in *R\** and to negative coupling between *Z* and *W*, with the latter effect being slightly stronger than the former.

If YFP abundance captures the production capacity Z of individual cells, and if fluctuations induced by the misP tag 𝑊 depend on the capacity to degrade proteins, our results imply that cells with a higher production capacity have a higher degradation capacity. The correlation coefficient between *Z* and *W* can, in fact, be computed from measurements of *G*, *R* and *R\** (Equation 18, Methods). We find *R(Z,W)* ≅-0.54, confirming that *Z* and *W* are anti-correlated.

### Numeric simulations confirm the coupling between protein production and degradation in single cells

To confirm that coupling between production and degradation leads to decoupling between passively- degraded (YFP) and actively degraded proteins (RFP-misP) we simulated the stochastic variability of proteins using the Gillespie algorithm with rate constants determined to yield average mRNA and protein copy numbers matching those of the literature (Fig 4A, Methods). Each simulated “yeast cell” consisted of a unique set of rate constants shared by all four proteins, except for protein degradation rates. Those were always identical for equivalent proteins but differed between passively (YFP and RFP) and actively degraded variants (YFP-misP, RFP-misP). The expression of all four proteins was simulated for 80 hours to reach equilibrium, at the end of which protein abundance was recorded for the four proteins. Finally, one thousand of these simulations were performed to obtain cell population statistics. We implemented two models of protein degradation. First, degradation rates were normally distributed across cells but were identical for YFP-misP and RFP-misP within cells. This simulates extrinsic noise in protein degradation. As expected, such cell-specific degradation rates reproduced the correlation between RFP-misP and YFP-misP observed experimentally (Fig 4B, model 1, R=0.8). However, the noise added by degradation did not decrease the correlation between YFP and RFP-misP to the extent observed experimentally (*R_sim_(G,R*)*=0.65). In a second model, the rate of protein degradation (*k_5_*) was coupled to the rate of production (arbitrarily chosen as *k_2_*). Equation 18 (Methods) enabled us to calculate the correlation that should exist between those rates to recapitulate the experimental results. We thus sampled values of *k5* such that, on average, the correlation between *log(k_2_)* and *log(k_5_)* would be 0.54. With this added constraint, the correlation between YFP-misP and RFP-misP remained high, with *R_sim_(G*,R*)*=0.72, but the correlation between YFP and RFP-misP decreased to *R_sim_(G,R*)* = 0.28, in good agreement with our experimental results where *R_exp_(G,R*)* = 0.32.

**Fig 4.**
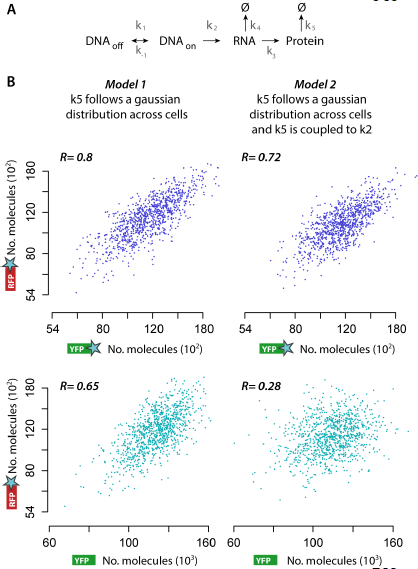
Stochastic simulations confirm that degradation capacity is coupled to production capacity in single cells. **(A)** Kinetic model used for the numeric simulations. Protein homeostasis was dependent on six parameters: the rate of DNA opening (k_1_) and closing (k_-1_), transcription from opened DNA (k_2_), mRNA translation (k_3_), mRNA degradation (k_4_) and protein degradation (k_5_). The parameters were identical in both models except for k_5_. In model 1, k_5_ was chosen randomly in each cell and stayed constant during simulations, subjecting YFP and RFP to identical degradation rates within cells, but to different rates across cells. This simulates extrinsic noise in protein degradation. Model 2 is identical to model 1, with the added constraint that k_5_ correlates with production k_2_. **(B)** Results of the numerical simulations for both models. In model 1, the abundance of YFP-misP and RFP-misP is correlated in single cells, as expected. However, YFP and RFP-misP remain correlated (R=0.65). In model 2, we couple protein production (arbitrarily chosen as k_2_) and degradation. Thus, the more YFP the higher the degradation rate, which decreased the correlation between YFP and RFP-misP while maintaining the correlation between RFP-misP and YFP-misP.

## Discussion and conclusions

### Functional implications of the findings

The fact that fluctuations in protein abundance can be coupled at the level of single cells brings about the question of function (4). The variability inherent to gene expression can be a constraint that is costly to suppress (57), but can also represent a beneficial, tunable and selectable trait as a primitive form of gene regulation (58), and in a bet-hedging context (31, 59–62). While bet-hedging is generally studied through variability of a single component, correlated fluctuations can be exploited to couple the fluctuations of many components together (63). The possibility to tune cell-to-cell fluctuations of multiple components could, for example, contribute to adjusting the stoichiometry of subunits in complexes at the level of single cells.

There may also be contexts where coupling is not desirable. For example, if preparing for all possible environmental stresses is too costly or hard to achieve functionally, cells may benefit from decoupling the expression of gene modules needed to overcome different types of stresses. Decoupling the expression of stress-response genes could allow individual cells to prepare to different kinds of stress, instead of all individuals preparing for all stresses. Doing so may increase the chance that at least a few cells survive sudden environmental stresses. In support of this conjecture, proteins involved in chemical homeostasis and defense response have high decay rates (64).

### Interaction between protein degradation and protein production

Modeling the experimental data suggested a connection between the rates of protein production and degradation (Fig 5). At the mRNA level, such dependence has been observed and is implemented through several mechanisms. For example, two RNA polymerase subunits (Rpb4 and Rpb7) were found to bind the transcribed mRNA to later direct it for degradation (65, 66). In another mechanism, a promoter element binding Rap1p stimulates both transcription and mRNA degradation (67). Additionally, RNA decay factors such as Xrn1 have been observed to enhance both the transcription and degradation of certain RNAs (68).

**Fig 5.**
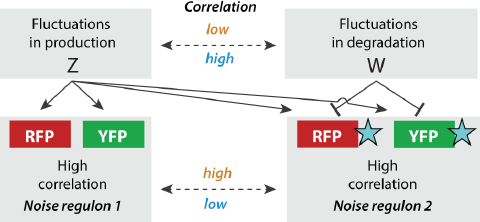
Protein degradation can decouple fluctuations in protein abundance. The expression of the fluorescent reporters is influenced by two sources of extrinsic noise, Z, and W, which capture effects associated with production and degradation respectively. The untagged RFP and YFP are both influenced by Z. The misP tag (blue star) couples RFP and YFP to an additional source of extrinsic noise, W, creating an incoherent feedforward loop. In the absence of coupling between Z and W, YFP and RFP-misP expression remain correlated. A positive coupling between Z and W results in RFP-misP being more degraded when YFP abundance is high, thus decreasing the correlation between YFP and RFP-misP in single cells.

Our data imply a similar linkage between production and degradation at the protein level. However, because untagged proteins are not subject to rapid degradation despite having the same promoter region as tagged proteins, the coupling between production and degradation is unlikely to involve a signal associated with the mRNA. The underlying mechanisms must thus be different from those described above. We hypothesize that the production-degradation linkage we observed reflects a more general mechanism. According to this hypothesis, a global extrinsic component – or, as coined by Stewart-Ornstein *et al.*, a “noise regulon” would be composed of proteins needed for growth, and include ribosome and metabolic enzymes, but also degradation factors. All the proteins in such a regulon would fluctuate together, thus simultaneously increasing production and degradation in a cell- specific manner. Such association may reflect a natural optimization of cells where, like in a factory, ramping up production naturally produces excess waste that needs to be cleared.

### Exploiting cellular noise may help characterize post-transcriptional regulation mechanisms

The strategy described in this work, whereby two fluorescent proteins differing in a specific feature capture the extrinsic noise component acting on that feature, is readily generalizable to dissect more regulatory mechanisms and pathways. The use of modified or synthetic proteins as “queries” could indeed reveal regulators of specific features. Yeast proteins that anti-correlate with RFP-misP could be candidate degradation factors. In contrast, yeast proteins that correlate with RFP-misP could be hypothesized as stabilizing chaperones or proteins subjected to the same degradation mechanism. We thus anticipate that cellular noise and co-fluctuating proteins will reveal mechanisms of regulation in biological systems, of and beyond transcriptional regulatory networks. To this aim, the framework introduced in this work, to analyze the extrinsic noise of non-equivalent reporters, will be instrumental.

## Materials and Methods

### Strains and plasmids

We employed two fluorescent proteins, Venus (YFP) and mCherry (RFP), which were cloned into plasmids suitable for genome integration at the *TRP1* locus (Supplementary Text 1). For genome integration, the plasmids were restricted by AccI (YFP) or BamHI (RFP), which released the cassette flanked by sequences bearing homology to the *TRP1* locus. The restricted fragment was transformed to BY4741 (YFP), or BY4742 (RFP) following an established protocol (69). Transformants were selected by antibiotic resistance (G418 for YFP, Hygromycin for RFP) and correct locus integration was verified by tryptophan auxotrophy. Mating was done by growth on solid YPD agar overnight followed by selection on synthetic media lacking methionine, lysine and supplemented with antibiotics selecting for the presence of both cassettes. The yeast strains used in this work are described in Tables S1 and S2.

### Microscope imaging

Cells were inoculated from their glycerol stock in 384-well glass-bottom optical plates (Matrical) with a pintool (FP1 pins, V&P Scientific) operated by a Tecan robot (Tecan Evo200 with MCA384 head). Cells were grown in YPD for a minimum of ten hours before they reached an optical density of at most 1, and were imaged. Imaging was performed with an automated Olympus microscope X83 coupled to a spinning disk confocal scanner (Yokogawa W1), using a 60X objective (Olympus, plan apo, 1.42 NA). Excitation was achieved with a green L.E.D for brightfield images, a 488 nm laser (Toptica, 100 mW) for YFP, and 561 nm laser (Obis, 75 mW) for RFP. Emission filter sets used to acquire the brightfield, YFP and RFP images were 520/28, 520/28 and 645/75 respectively. The same triple-band dichroic mirror was used for all channels (405/488/561, Yokogawa). Images were recorded on two Hamamatsu Flash4-V2 cameras, one for the brightfield and YFP channels and the second for the RFP channel. Each image set was composed of two brightfield (BF) images (one in focus and one defocused to facilitate cell segmentation, each with 50 ms exposure) as well as one image for each fluorescent channel (500 ms exposure for YFP and 700 ms exposure for RFP). The focus was maintained throughout the experiment by hardware autofocus (Olympus z-drift compensation system).

### Image analysis

Images were processed with ImageJ by custom algorithms. Individual cells were segmented from the brightfield images and statistics for all four images (in-focus brightfield, out-of- focus brightfield, YFP, and RFP) were recorded. Fluorescence intensity was estimated from the 30^th^ quantile of pixel intensity within each cell. All tabulated data were analyzed in R. Several filters were applied to the data extracted from the images (Text S3, Fig S6).

### Modeling reporters subject to a common noise source

Untagged fluorescent reporters are considered equivalent and share the same extrinsic noise *Z.* We model their expression using the framework introduced by Elowitz *et al.* (8), with

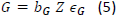

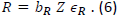

Here, *G* and *R* are single-cell fluorescence measured in the green (GFP or YFP) and red (RFP) channels respectively. *Z* represents the extrinsic noise, while 𝜖 models the noise intrinsic to the process of gene expression. The *b* constants account for differences in average protein abundances and differences in abundance-to-fluorescence scaling between the two reporters. We use a multiplicative model between sources of noise because protein abundance is the result of chemical reactions with mass action kinetics. With such kinetics, protein abundance is given by the product of kinetic rates. For example, consider a cell with 3-fold more transcription activity and 3-fold more translation activity for a particular gene when compared to average. In such a cell, the fold change at the protein level will be 9-fold compared to average, and not 6-fold. This aspect of the model is important when comparing and modeling the correlation between fluorescent reporters whose abundance we alter experimentally, as we do here.

Log-transforming the equations linearizes them. We apply x:=log(x) for all the variables in the model and redefine the 𝑏 constants such that 𝑍 and 𝜖 have mean 0 which yields

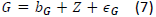

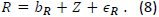

In this context, the total variance of single-cell fluorescence can be decomposed into contributions of extrinsic and intrinsic noise, with

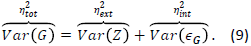

To determine how coupling reporters to different sources of noise alter the correlation between fluorescence, we note that reporters lacking the misP tag are only influenced by *Z*. In this simple case, the correlation between *G* and *R* is derived from Equations 7 and 8 as

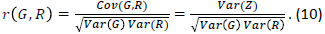

Here, the correlation *r(G,R)* quantifies the amount of extrinsic noise *Var(Z)* relative to the total noise of the fluorescent proteins. This formula, based on the correlation coefficient, is a normalized form of the original formula based on the covariance to estimate extrinsic noise (8).

### Modeling of reporters subjected to different sources of noise

The model of Equation 10 assumes the two fluorescent reporters to be subject to the same source of extrinsic noise *Z*. However, the misP tag subjects the fluorescent reporter *R\** to an additional source of extrinsic noise *W*. We model the effect of *W* on the abundance *R\** as

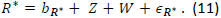

Coupling *R\** to an additional noise source *W* tends to increase fluctuations, as shown by computing *Var(R*)* as a function of *Var(R)*, which gives

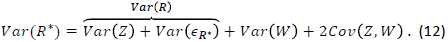

Since *Var(W)* > 0, coupling *R\** to *W* tends to increase the variance of *R\**, unless *Z* and *W* are strongly anti-coupled, *i.e.*, *Cov(Z,W)* << 0. Using Equations 7 and 11 which define *G* and *R\**, we can derive how *Z* and *W* impact the correlation between *G* and *R\**,

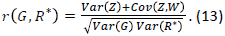

By dividing this equation by Equation 10, we obtain an expression for how *Z* and *W* impact the correlation between the two pairs of reporters,

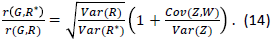

This equation suggests two mechanisms through which the misP tag can alter the correlation between the two reporters. In one mechanism, *W* injects more noise into *R\**. This increase *Var(R*)* (first term), and thus decreases *r(G,R*)* relative to *r(G,R)*. In another mechanism, *W* is anti-coupled to *Z* such that fluctuations in *Z* are partially canceled by anti-fluctuations in *W*. In this scenario, a negative covariance between *Z* and *W* decreases *r(G,R*)*.

To quantify the strength of the coupling between *Z* and *W*, we compute *r(Z,W)*. We first use Equations 7 and 8, which define *G* and *R* to find

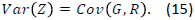

From Equation 11 which defines *R\** and its equivalent form for *G\**, we find the variance of *W* as

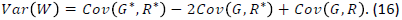

By computing *Cov(G,R)* and *Cov(G,R*)* and solving for *Cov(Z,W)*, we can show that

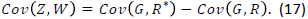

Lastly, combining Equations 15, 16 and 17, we obtain

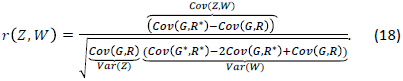

### Numerical simulations

The simulations were based on the Gillespie algorithm (70) adapted from Bahar- Halpern *et al*. (71). The algorithm was modified to account for protein translation and degradation, and was ported to the R language. The rates used in the simulations were for gene opening (k_1_=5/h) and closing (k_-1_=5/h) (72), mRNA transcription (k_2_~N(20/h,0.12/h^2^) and degradation (k_3_=0.5/h), protein translation (k_4_~N(30/h,0.05/h^2^)) and protein degradation (k_5_=0.005/h for untagged YFP or RFP). These rates gave copy numbers of mRNA (average of 20 per cell) and proteins (average of 120,000 per cell) comparable to expected values (73, 74) (BNID 104745,104185). We simulated the impact of the misP tag using two models of protein degradation. In model 1, values of k_5_ ~ N(0.05/h,0.006/h^2^) were identical for YFP-misP and RFP-misP in single cells. Model 2 was identical to model 1, with the added constraint that values of k_5_ were sampled so that log(k2) and log(k5) would show a correlation expected to be 0.54.

## Statistical analysis

Two-tailed Welch exact t-test was used to compare mean values of measurements series. All correlations in this works were calculated with the Pearson coefficient. The Fisher exact test was used to evaluate the significance of the correlations.

## Acknowledgments

We thank Steve Altschuler, Naama Barkai, Michael Elowitz, and Eran Segal for valuable discussions and comments. We thank Naama Barkai, Hagen Hofmann, Amnon Horovitz, Shalev Itzkovitz and Schraga Schwartz for insightful comments on the manuscript. We thank Maya Schuldiner for kindly providing the BY4741/2 strains.

## Author contributions

OM and EDL designed the study. OM performed the experiments. AS and EDL processed the images. ES designed the yeast tagging plasmid. JH, OM, and EDL developed the theoretical analysis framework. OM, JH, and EDL analyzed the data. OM, JH, and EDL wrote the manuscript with comments from all authors. All authors read and approved the final manuscript.

## References

1. Raser MJ, O’Shea KE. Noise in gene expression: origins, consequences, and control. Science. 2005;309:2010–3.

2. Newman RJ, Ghaemmaghami S, Ihmels J, Breslow KD, Noble M, DeRisi LJ, et al. Single-cell proteomic analysis of S. cerevisiae reveals the architecture of biological noise. Nature. 2006;441:840–6.

3. Raj A, van Oudenaarden A. Nature, nurture, or chance: stochastic gene expression and its consequences. Cell. 2008;135(2):216–26.

4. Eldar A, Elowitz MB. Functional roles for noise in genetic circuits. Nature 2010;467(7312):167–73.

5. Raser JM, O’Shea EK. Control of stochasticity in eukaryotic gene expression. Science. 2004;304(5678):1811–4.

6. Bar-Even A, Paulsson J, Maheshri N, Carmi M, O’Shea E, Pilpel Y, et al. Noise in protein expression scales with natural protein abundance. Nature genetics 2006;38:636–43.

7. Taniguchi Y, Choi PJ, Li GW, Chen H, Babu M, Hearn J, et al. Quantifying E. coli proteome and transcriptome with single-molecule sensitivity in single cells. Science. 2010;329(5991):533–8

8. Elowitz BM, Levine JA, Siggia DE, Swain SP. Stochastic gene expression in a single cell. Science 2002;297:1183–6.

9. Swain SP, Elowitz BM, Siggia DE. Intrinsic and extrinsic contributions to stochasticity in gene expression. Proceedings of the National Academy of Sciences of the United States of America 2002;99:12795–800.

10. Bowsher CG, Swain PS. Identifying sources of variation and the flow of information in biochemical networks. Proceedings of the National Academy of Sciences of the United States of America. 2012;109(20):E1320–E8.

11. Hilfinger A, Paulsson J. Separating intrinsic from extrinsic fluctuations in dynamic biological systems. Proc Natl Acad Sci U S A 2011;108(29):12167–72.

12. Padovan-Merhar O, Nair GP, Biaesch AG, Mayer A, Scarfone S, Foley SW, et al. Single mammalian cells compensate for differences in cellular volume and DNA copy number through independent global transcriptional mechanisms. Mol Cell 2015;58(2):339–52.

13. Battich N, Stoeger T, Pelkmans L. Control of Transcript Variability in Single Mammalian Cells. Cell 2015;163(7):1596–610.

14. Colman-Lerner A, Gordon A, Serra E, Chin T, Resnekov O, Endy D, et al. Regulated cell-to-cell variation in a cell-fate decision system. Nature. 2005;437(7059):699–706.

15. Volfson D, Marciniak J, Blake JW, Ostroff N, Tsimring SL, Hasty J. Origins of extrinsic variability in eukaryotic gene expression. Nature 2006;439:861–4.

16. Jaitin DA, Kenigsberg E, Keren-Shaul H, Elefant N, Paul F, Zaretsky I, et al. Massively parallel single-cell RNA-seq for marker-free decomposition of tissues into cell types. Science. 2014;343(6172):776–9. Epub 2014/02/18.

17. Setty M, Tadmor MD, Reich-Zeliger S, Angel O, Salame TM, Kathail P, et al. Wishbone identifies bifurcating developmental trajectories from single-cell data. Nat Biotechnol 2016;34(6):637–45. Epub 2016/05/03.

18. Zeisel A, Munoz-Manchado AB, Codeluppi S, Lonnerberg P, La Manno G, Jureus A, et al. Brain structure. Cell types in the mouse cortex and hippocampus revealed by single-cell RNA- seq. Science. 2015;347(6226):1138–42. Epub 2015/02/24.

19. Klein AM, Mazutis L, Akartuna I, Tallapragada N, Veres A, Li V, et al. Droplet barcoding for single-cell transcriptomics applied to embryonic stem cells. Cell 2015;161(5):1187–201. Epub 2015/05/23.

20. Pedraza JM, van Oudenaarden A. Noise propagation in gene networks. Science. 2005;307(5717):1965–9.

21. Stewart-Ornstein J, Weissman SJ, El-Samad H. Cellular noise regulons underlie fluctuations in Saccharomyces cerevisiae. Molecular cell 2012;45:483–93.

22. Sigal A, Milo R, Cohen A, Geva-Zatorsky N, Klein Y, Liron Y, et al. Variability and memory of protein levels in human cells. Nature 2006;444(7119):643–6.

23. Dunlop MJ, Cox RS, 3rd, Levine JH, Murray RM, Elowitz MB. Regulatory activity revealed by dynamic correlations in gene expression noise. Nat Genet. 2008;40(12):1493–8.

24. Cox CD, McCollum JM, Allen MS, Dar RD, Simpson ML. Using noise to probe and characterize gene circuits. Proceedings of the National Academy of Sciences of the United States of America 2008;105(31):10809–14.

25. Snijder B, Pelkmans L. Origins of regulated cell-to-cell variability. Nature Reviews Molecular Cell Biology 2011;12(2):119–25.

26. Bhosale R, Jewell BJ, Hollunder J, Koo KAJ, Vuylsteke M, Michoel T, et al. Predicting gene function from uncontrolled expression variation among individual wild-type Arabidopsis plants. The Plant cell 2013;25:2865–77.

27. Geiler-Samerotte AK, Bauer RC, Li S, Ziv N, Gresham D, Siegal LM. The details in the distributions: why and how to study phenotypic variability. Current Opinion in Biotechnology 2013;24:752–9.

28. Padovan-Merhar O, Raj A. Using variability in gene expression as a tool for studying gene regulation. Wiley interdisciplinary reviews Systems biology and medicine 2013;5:751–9.

29. Farkash-Amar S, Zimmer A, Eden E, Cohen A, Geva-Zatorsky N, Cohen L, et al. Noise genetics: inferring protein function by correlating phenotype with protein levels and localization in individual human cells. PLoS genetics. 2014;10.

30. Lipinski-Kruszka J, Stewart-Ornstein J, Chevalier WM, El-Samad H. Using dynamic noise propagation to infer causal regulatory relationships in biochemical networks. ACS synthetic biology 2015;4:258–64.

31. Levy SF, Ziv N, Siegal ML. Bet Hedging in Yeast by Heterogeneous, Age-Correlated Expression of a Stress Protectant. Plos Biology. 2012;10(5).

32. Rhee A, Cheong R, Levchenko A. Noise decomposition of intracellular biochemical signaling networks using nonequivalent reporters. Proc Natl Acad Sci U S A 2014;111(48):17330–5. Epub 2014/11/19.

33. Conlon P, Gelin-Licht R, Ganesan A, Zhang J, Levchenko A. Single-cell dynamics and variability of MAPK activity in a yeast differentiation pathway. Proc Natl Acad Sci U S A. 2016;113(40):E5896–E905. Epub 2016/09/22.

34. Thattai M, van Oudenaarden A. Intrinsic noise in gene regulatory networks. Proceedings of the National Academy of Sciences of the United States of America. 2001;98:8614–9.

35. Blake WJ, M ka, Cantor CR, Collins JJ. Noise in eukaryotic gene expression. Nature 2003;422(6932):633–7.

36. Weinberger SL, Burnett CJ, Toettcher EJ, Arkin PA, Schaffer VD. Stochastic gene expression in a lentiviral positive-feedback loop: HIV-1 Tat fluctuations drive phenotypic diversity. Cell 2005;122:169–82.

37. Maamar Hd, Raj A, Dubnau D. Noise in gene expression determines cell fate in Bacillus subtilis. Science: American Association for the Advancement of Science; 2007. p. 526–9.

38. Cai L, Dalal CK, Elowitz MB. Frequency-modulated nuclear localization bursts coordinate gene regulation. Nature. 2008;455(7212):485–U16.

39. Kalmar T, Lim C, Hayward P, Muoz-Descalzo S, Nichols J, Garcia-Ojalvo J, et al. Regulated fluctuations in nanog expression mediate cell fate decisions in embryonic stem cells. PLoS biology. 2009;7.

40. Yang S, Kim S, Rim Lim Y, Kim C, An JH, Kim J-H, et al. Contribution of RNA polymerase concentration variation to protein expression noise. Nature communications. 2014;5.

41. Sherman MS, Lorenz K, Lanier MH, Cohen BA. Cell-to-Cell Variability in the Propensity to Transcribe Explains Correlated Fluctuations in Gene Expression. Cell Systems. 2015;1.

42. Haque A, Engel J, Teichmann SA, Lonnberg T. A practical guide to single-cell RNA- sequencing for biomedical research and clinical applications. Genome medicine. 2017;9(1):75. Epub 2017/08/20.

43. Yuan GC, Cai L, Elowitz M, Enver T, Fan G, Guo G, et al. Challenges and emerging directions in single-cell analysis. Genome biology. 2017;18(1):84. Epub 2017/05/10.

44. Vogel C, Marcotte EM. Insights into the regulation of protein abundance from proteomic and transcriptomic analyses. Nat Rev Genet 2012;13(4):227–32.

45. Liu Y, Beyer A, Aebersold R. On the Dependency of Cellular Protein Levels on mRNA Abundance. Cell 2016;165(3):535–50.

46. Albayrak C, Jordi CA, Zechner C, Lin J, Bichsel CA, Khammash M, et al. Digital Quantification of Proteins and mRNA in Single Mammalian Cells. Mol Cell 2016;61(6):914–24. Epub 2016/03/19.

47. Yofe I, Weill U, Meurer M, Chuartzman S, Zalckvar E, Goldman O, et al. One library to make them all: streamlining the creation of yeast libraries via a SWAp-Tag strategy. Nat Methods 2016;13(4):371–8. Epub 2016/03/02.

48. Evans T, Rosenthal ET, Youngblom J, Distel D, Hunt T. Cyclin: a protein specified by maternal mRNA in sea urchin eggs that is destroyed at each cleavage division. Cell 1983;33(2):389–96.

49. Belle A, Tanay A, Bitincka L, Shamir R, O’Shea EK. Quantification of protein half-lives in the budding yeast proteome. Proceedings of the National Academy of Sciences of the United States of America. 2006;103(35):13004–9.

50. Gsponer Jr, Futschik EM, Teichmann AS, Babu MM. Tight regulation of unstructured proteins: from transcript synthesis to protein degradation. Science 2008;322:1365–8.

51. Kristensen AR, Gsponer J, Foster LJ. Protein synthesis rate is the predominant regulator of protein expression during differentiation. Mol Syst Biol. 2013;9.

52. dos Reis M, Savva R, Wernisch L. Solving the riddle of codon usage preferences: a test for translational selection. Nucleic Acids Res 2004;32(17):5036–44.

53. Remy I, Michnick SW. Clonal selection and in vivo quantitation of protein interactions with protein-fragment complementation assays (vol 96, pg 5394, 1999). Proceedings of the National Academy of Sciences of the United States of America. 1999;96(13):7610-.

54. Hipp MS, Park SH, Hartl FU. Proteostasis impairment in protein-misfolding and - aggregation diseases. Trends Cell Biol 2014;24(9):506–14.

55. Stansfield I, Jones KM, Herbert P, Lewendon A, Shaw WV, Tuite MF. Missense translation errors in Saccharomyces cerevisiae. J Mol Biol 1998;282(1):13–24.

56. Tyedmers J, Mogk A, Bukau B. Cellular strategies for controlling protein aggregation. Nature reviews Molecular cell biology 2010;11(11):777–88. Epub 2010/10/15.

57. Lestas I, Vinnicombe G, Paulsson J. Fundamental limits on the suppression of molecular fluctuations. Nature 2010;467(7312):174–8.

58. Wolf L, Silander OK, van Nimwegen E. Expression noise facilitates the evolution of gene regulation. eLife. 2015;4. Epub 2015/06/18.

59. Kussell E, Leibler S. Phenotypic diversity, population growth, and information in fluctuating environments. Science 2005;309(5743):2075–8.

60. Wolf DM, Vazirani VV, Arkin AP. Diversity in times of adversity: probabilistic strategies in microbial survival games. J Theor Biol 2005;234(2):227–53.

61. Acar M, Mettetal JT, van Oudenaarden A. Stochastic switching as a survival strategy in fluctuating environments. Nature Genetics. 2008;40(4):471–5.

62. Frankel NW, Pontius W, Dufour YS, Long J, Hernandez-Nunez L, Emonet T. Adaptability of non-genetic diversity in bacterial chemotaxis. eLife. 2014;3. Epub 2014/10/04.

63. Garcia-Bernardo J, Dunlop MJ. Noise and Low-Level Dynamics Can Coordinate Multicomponent Bet Hedging Mechanisms. Biophysical Journal 2015;108:184–93.

64. Schwanhausser B, Busse D, Li N, Dittmar G, Schuchhardt J, Wolf J, et al. Global quantification of mammalian gene expression control. Nature 2011;473(7347):337–42. Epub 2011/05/20.

65. Dori-Bachash M, Shalem O, Manor YS, Pilpel Y, Tirosh I. Widespread promoter-mediated coordination of transcription and mRNA degradation. Genome biology. 2012;13(12):R114. Epub 2012/12/15.

66. Shalem O, Groisman B, Choder M, Dahan O, Pilpel Y. Transcriptome Kinetics Is Governed by a Genome-Wide Coupling of mRNA Production and Degradation: A Role for RNA Pol II. Plos Genetics. 2011;7(9).

67. Bregman A, Avraham-Kelbert M, Barkai O, Duek L, Guterman A, Choder M. Promoter elements regulate cytoplasmic mRNA decay. Cell 2011;147(7):1473–83. Epub 2011/12/27.

68. Haimovich G, Medina DA, Causse SZ, Garber M, Millan-Zambrano G, Barkai O, et al. Gene Expression Is Circular: Factors for mRNA Degradation Also Foster mRNA Synthesis. Cell 2013;153(5):1000–11.

69. Knop M, Siegers K, Pereira G, Zachariae W, Winsor B, Nasmyth K, et al. Epitope tagging of yeast genes using a PCR-based strategy: more tags and improved practical routines. Yeast. 1999;15(10B):963–72.

70. Gillespie DT. Exact Stochastic Simulation of Coupled Chemical-Reactions. J Phys Chem-Us 1977;81(25):2340–61.

71. Bahar Halpern K, Caspi I, Lemze D, Levy M, Landen S, Elinav E, et al. Nuclear Retention of mRNA in Mammalian Tissues. Cell Rep 2015;13(12):2653–62. Epub 2015/12/30.

72. Zenklusen D, Larson DR, Singer RH. Single-RNA counting reveals alternative modes of gene expression in yeast. Nature structural & molecular biology 2008;15(12):1263–71. Epub 2008/11/18.

73. Milo R, Jorgensen P, Moran U, Weber G, Springer M. BioNumbers—the database of key numbers in molecular and cell biology. Nucleic Acids Res. 2010;38(Database issue):D750–3. Epub 2009/10/27.

74. Wang M, Herrmann CJ, Simonovic M, Szklarczyk D, von Mering C. Version 4.0 of PaxDb: Protein abundance data, integrated across model organisms, tissues, and cell-lines. Proteomics. 2015;15(18):3163–8. Epub 2015/02/07.

